# Open channel droplet-based microfluidics

**DOI:** 10.1101/436675

**Authors:** Samuel B. Berry, Jing J. Lee, Jean Berthier, Erwin Berthier, Ashleigh B. Theberge

## Abstract

Droplet-based microfluidics enables compartmentalization and controlled manipulation of small volumes. Open microfluidics provides increased accessibility, adaptability, and ease of manufacturing compared to closed microfluidic platforms. Here, we begin to build a toolbox for the emerging field of open channel droplet-based microfluidics, combining the ease of use associated with open microfluidic platforms with the benefits of compartmentalization afforded by droplet-based microfluidics. We develop fundamental microfluidic features to control droplets flowing in an immiscible carrier fluid within open microfluidic systems. Our systems use capillary flow to move droplets and carrier fluid through open channels and are easily fabricated through 3D printing, micromilling, or injection molding; further, droplet generation can be accomplished by simply pipetting an aqueous droplet into an empty open channel. We demonstrate on-chip incubation of multiple droplets within an open channel and subsequent transport (using an immiscible carrier phase) for downstream experimentation. We also present a method for tunable droplet splitting in open channels driven by capillary flow. Additional future applications of our toolbox for droplet manipulation in open channels include cell culture and analysis, on-chip microscale reactions, and reagent delivery.

## Introduction

Open microfluidic systems offer many advantages for conducting life science experimentation including pipette accessibility, simple fabrication techniques with biocompatible materials, independence from pumps and external flow generators, and customizability.^1^ Here, we describe a biphasic system driven by capillary forces that enables the control and manipulation of multiple droplets within an open channel devoid of any electrical or pneumatic actuation systems, in a fully open, pipette-accessible platform. We demonstrate a new open channel system for prolonged static droplet incubation in channel followed by capillary-driven translation of discrete droplets for downstream analysis, as well as tunable droplet splitting in open channels.

Droplet-based microfluidics advances the capabilities of traditional single-phase microfluidic platforms through compartmentalization of reaction components into discrete micro- to picoliter volumes, enabling decreased reagent consumption and use of valuable, low-volume samples that may otherwise be expensive or difficult to obtain.^2,3^ Translation of assays to droplet-based platforms allows users to precisely form, manipulate, and transport small volumes for use in cell-based assays, chemical synthesis, and biochemical analyses.^2^ Droplet-based systems for an extensive range of functions have been described^3^, and droplet manipulation methods such as incubation^4,5^, reagent addition^6^, and splitting^7,8^ have been developed. However, most current droplet-based microfluidic approaches rely on complex designs and multistep fabrication methods (e.g., photolithography, bonding) to create closed-channel platforms and often use external pumps and actuators to manipulate flow, allowing them to perform specialized functions but limiting their wide-spread adoption beyond engineering and physical science laboratories.^3^ Recent work by Li et al.^9^ overcomes some of the fabrication challenges of traditional droplet systems by using an open paper-based device, but the flow still requires external syringe pumps.

Systems utilizing open fluidic channels (e.g., channels devoid of a ceiling, devoid of a ceiling and floor, or devoid of lateral walls) and surface tension driven flow have emerged as alternatives to closed channel, pump-driven microfluidic platforms due to their relative ease of design, fabrication, and use.^1,10^ Open channel platforms do not require bonding and can be fabricated in a single step using micromilling^11-13^ or high-volume fabrication techniques such as injection molding.^14,15^ Open platforms provide improved accessibility (e.g., pipette, automated reagent delivery systems) to users to manipulate experimental conditions through direct addition or removal of reagents at any point on the platform.^16^ Additionally, open channel systems can be driven by capillary flow in a manner similar to that of closed capillary systems. Capillary flow removes the need for external flow drivers and improves the robustness and functionality of the platform, as the mechanism for flow is built into the device.^10^ Recently, we presented an analytical model, numerical simulations, and experimental validation that described the behavioral modes of a single immiscible droplet placed in an open channel where a carrier flow occurs^17^; we found that an immiscible droplet can behave in a number of fundamentally different ways (remain static in the channel, translate at the leading edge of the carrier fluid, or detach from the walls of the channel and flow with the carrier fluid).

In our prior work^17^, we also showed that multiple aqueous droplets can be created and transported by pipetting the aqueous phase into an oil carrier phase that is already flowing through the device based on capillary flow. In the present manuscript, we developed a new capability, which enables extended incubation of droplets within the channel in the absence of the carrier phase, followed by introduction of the carrier phase in the channel using spontaneous capillary flow, and subsequent movement of the droplets. In contrast to our prior work, the present manuscript enables longer residence times of the droplets within the channels since they can be incubated for multiple hours before the carrier phase is added. Pipetting multiple aqueous droplets directly into an empty channel, incubating them, and *then* translating the train of droplets using a capillary-driven immiscible phase presents further challenges, as conditions such as surface wetting, evaporation, droplet merging, and satellite droplet formation all must be accounted for.

Here, we build a toolbox of droplet manipulation capabilities for open channel droplet-based microfluidics. We describe new open channel systems in which multiple discrete droplets can be placed into an open channel, incubated *in situ*, and then translated downstream either with droplet merging or without droplet merging, depending on the desired application. We also demonstrate an open microfluidic droplet splitting method to enable a parent droplet to be aliquoted into tailored smaller droplets (equal or unequal volumes) for multiplexed processing and readouts. Our open microfluidic systems rely on the surface interactions between the aqueous droplets, organic carrier phase, and channel surface which can be altered to fit various experimental needs; additionally, reliance on capillary-driven flow in an open channel removes the need for flow-generation devices and enables direct user access to the system at any time point. These platform functionalities (i.e., merging/splitting, incubation, user access) can help streamline large and cumbersome screening experiments that rely on manual pipetting, mixing, and splitting for sample generation, where manual processing can negatively contribute to assay time, sample loss, and costs associated with instrument usage. In future applications, these functionalities can be adapted and applied to array generation, sample preparation, and multiplexing.

## Materials and Methods

### Materials

Droplets were created with deionized (DI) water (Type II, Harleco; Fisher Scientific, Hampton, NH) tinted with either yellow or green dye (Spice Supreme; Gel Spice Company, Bayonne, NJ) at a concentration of 10% or 1% (v/v), respectively. The carrier fluids were: toluene (Fisher Scientific, Figure 1) or 1-pentanol (Acros Organics, Thermo Fisher Scientific, Waltham, MA, Figures 1, 2, 3, and 4). All carrier fluids solvents were tinted with Solvent Green 3 (Sigma-Aldrich, St. Louis, MO) at a concentration of 0.50 mg/mL.

**Figure 1.**
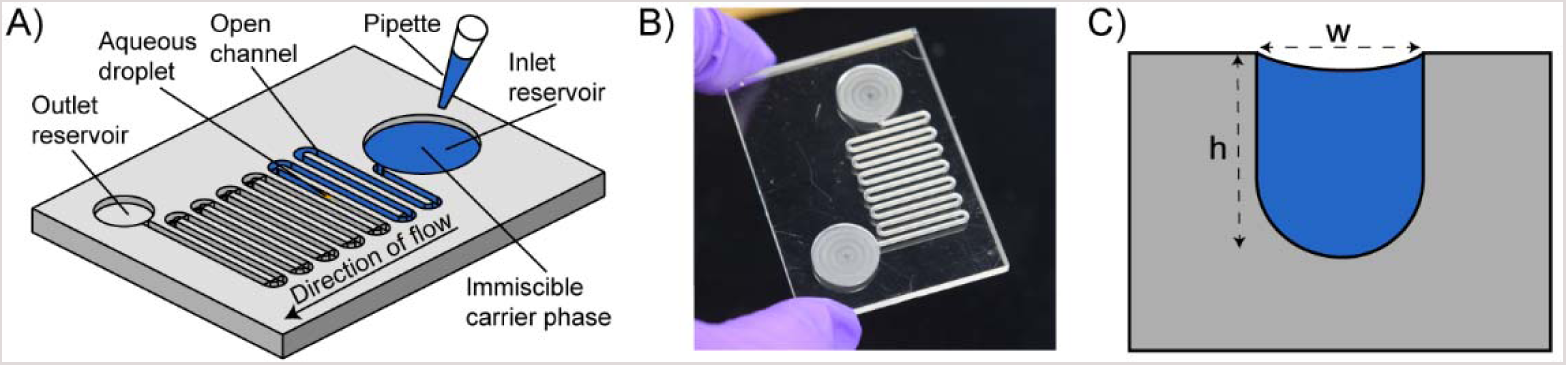
General platform design and modes of operation for translating aqueous droplets via capillary flow of an organic carrier phase. A) Schematic representation of open channel platform illustrating addition of organic carrier fluid (blue) with aqueous droplets (yellow) present in the channel; B) image of open channel platform; C) cross sectional schematic of channel (w = 0.90 mm and h = 1.0 mm).

**Figure 2.**
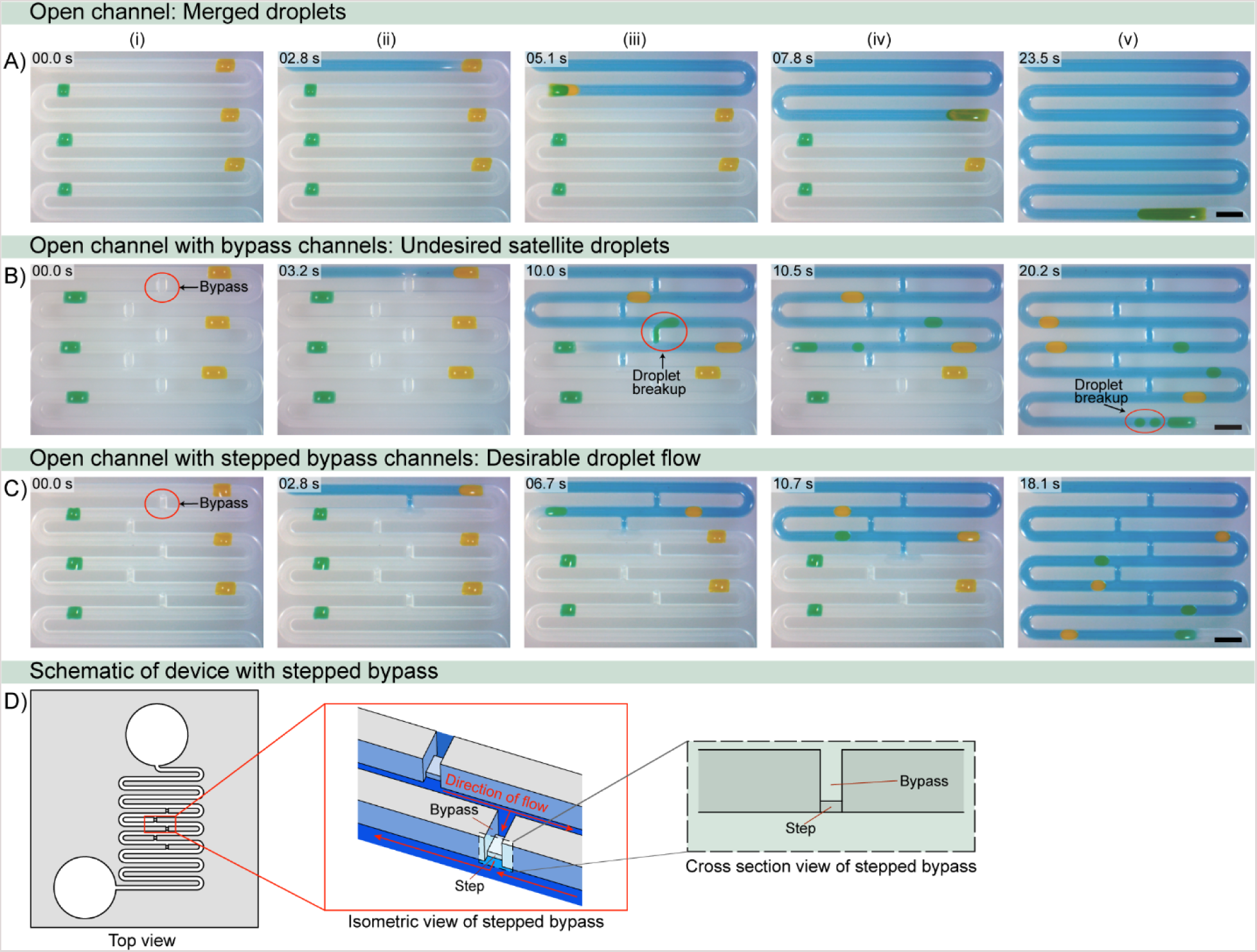
Open channel droplet incubation and transport. Droplets (alternating yellow and green for visualization) are incubated in an open channel without (A) or with (B-C) bypass channels (red circles); i-v) carrier fluid (blue) is pipetted in the inlet and flows down the channel via capillary flow, translating the droplets down the channel to the outlet reservoir. Bypass channels (B-C) prevent coalescence of preincubated droplets by inserting immiscible carrier fluid between aqueous droplets as they flow downstream. C) Incorporation of stepped bypass improves flow in the main channel and prevents formation of satellite droplets and droplet stagnation. D) Schematic of stepped bypass showing an isometric and cross-sectional view of the step in the bypass. Scale bar: 2 mm. Timestamps correspond to the addition of the carrier fluid (0.0 s) and not total droplet incubation time. Videos for A-C are included in the SI (Videos S3-S5).

**Figure 3.**
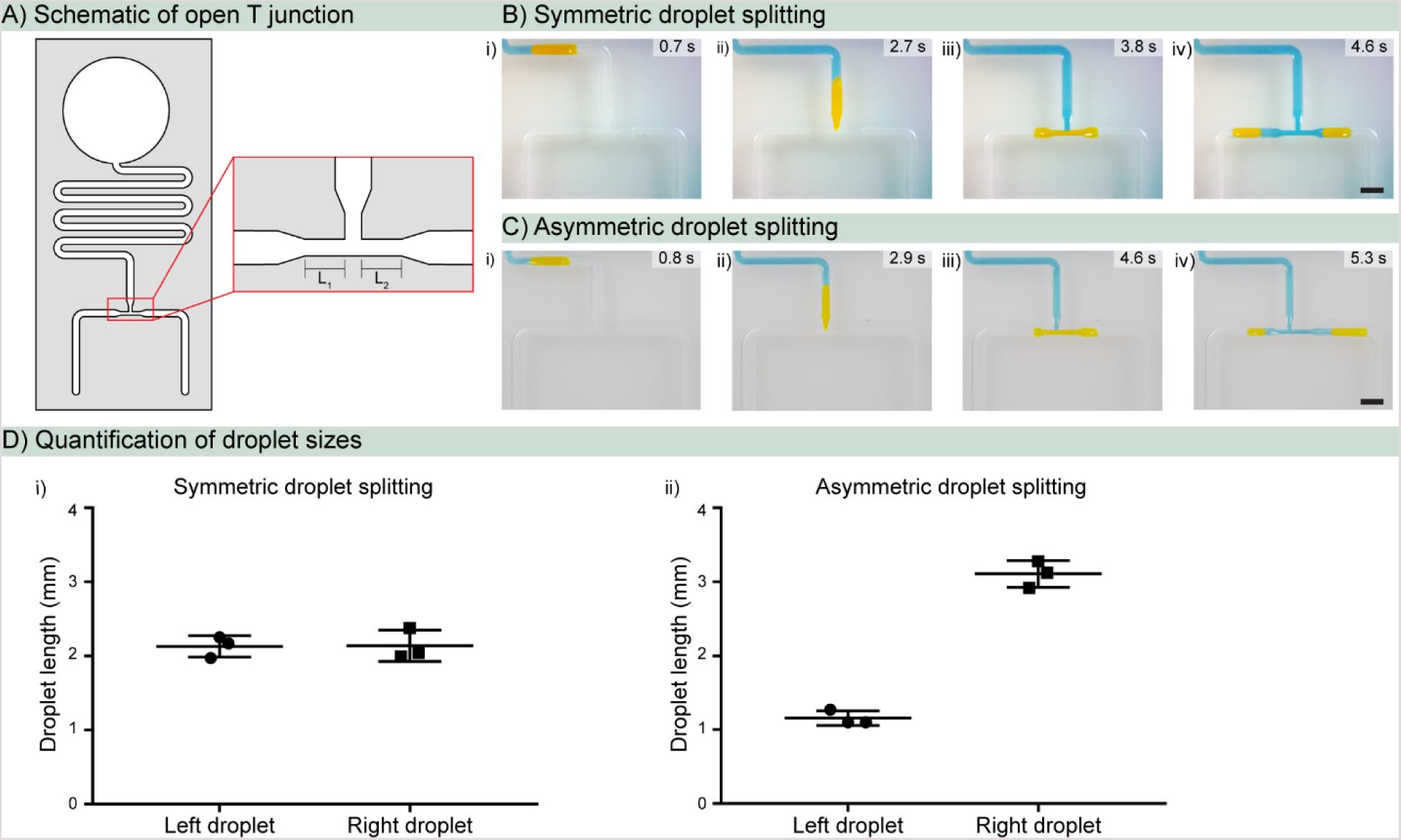
Controlled and adjustable droplet splitting in open channels with SCF. A) Schematic of T junction showing branches (L_1_ and L_2_) within the junction; B-C) i-ii) a droplet in the channel above a T junction with symmetric (B) or asymmetric (C) branch lengths is translated toward the junction via SCF; iii) the droplet fills both branches in the junction and slows upon reaching the channel expansion after the junction due to temporary pinning; iv) the droplet splits into two discrete droplets dependent upon the branch length; D) quantification of daughter droplet sizes after splitting in T junctions with symmetric (i) or asymmetric (ii) branch ratios (the three points plotted are from three different devices; mean and standard deviation are indicated). Scale bar: 2 mm. Videos for B and C are included in the SI (Videos S6 and S7).

**Figure 4.**
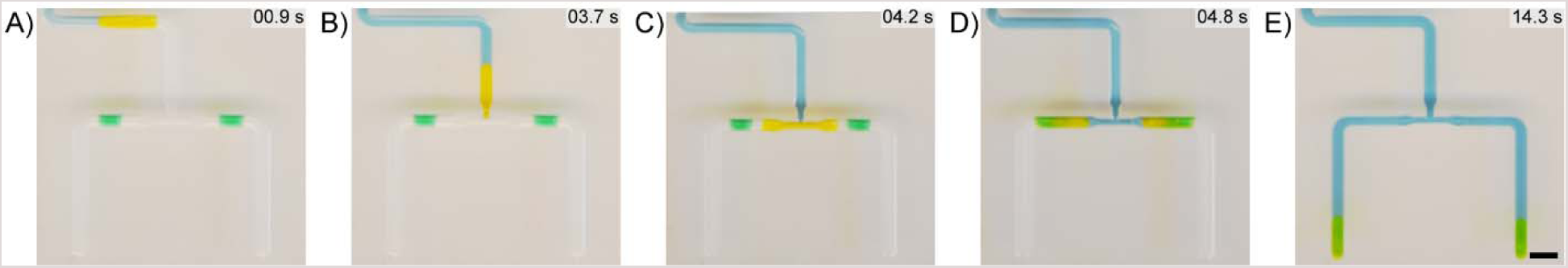
Workflow for droplet splitting and merging with downstream droplets to model reagent delivery. A yellow droplet (representing a primary reagent) was pipetted above the T junction while green droplets (representing secondary reagents) were added after the junction. A-B) Carrier fluid translates the yellow droplet via SCF into the junction; C) the droplet splits equally and is delivered (D) to the secondary reagents in the channel; E) the droplet and reagent mix as the droplet flows down the channel. Scale bar: 2 mm. Corresponding video is included in the SI (Video S8).

### Device fabrication

Devices were designed with Solidworks 2017 (Solidworks, Waltham, MA) and converted to .TAP files with SprutCam 10 (SprutCam, Naberezhnye Chelny, Russia). The devices were milled on poly(methyl methacrylate) (PMMA) sheets of 3.175 mm thickness (McMaster-Carr, Santa Fe Springs, CA) using a Tormach PCNC 770 mill (Tormach, Waunakee, WI). All device channels were milled with ball endmills (Performance Micro Tool, Janesville, WI) with a cutter diameter of 1/32” (TR-2-0313-BN) or 1/64” (TR-2-0150-BN) to create round-bottom channels. After milling, the devices were rinsed with DI water, sonicated in 70% (v/v) ethanol, and rinsed again with DI water. The devices were then dried with compressed air prior to use.

### Device design and testing

The main device dimensions are a channel width of 0.90 mm and a channel depth of 1.0 mm, with smaller channels included for carrier fluid bypass (0.5 mm wide, 1.0 mm length, 0.2 mm step height) (Figure 2D) and splitting (1.0 mm branch length, 0.45 mm channel width); the detailed dimensions of the devices and features are included in the SI (Figure S2). Computer aided design (CAD) files (.STEP) are also included in the SI. Aqueous droplets with a volume of 1.0 μL (Figures 1 and 2) or 3.0 μL (Figures 3 and 4) were generated in the channel with a pipette. Carrier fluids with a volume of 240 μL were dispensed in the inlet reservoir of the channel. Droplets were imaged and analyzed with ImageJ (National Institutes of Health, MD) for quantification (Figure 3 and S3). To prevent evaporation, devices were placed inside of a humidified Nunc™ Omnitray™ (Thermo Fisher, Frederick, MD) surrounded by 1.5 mL of sacrificial water droplets (≈ 50 μL/droplet), and the Omnitray was then placed inside a secondary humidified bioassay dish (#240835, Thermo Fisher) containing 100 mL of sacrificial water for extended incuabtions.

### Imaging

Images and videos were acquired using a MU1403B High Speed Microscope Camera mounted on an Amscope SM-3TZ-80S stereoscope (Amscope, Irvine, CA) unless otherwise noted. For Figure 1B, 3C and 4, images and videos were obtained with a Nikon-D5300 ultra-high resolution SLR camera (Tokyo, Japan).

## Results and Discussion

### Capillary-driven flow of droplets in an open channel

While the dynamics and behavior of single phase capillary flow within an open system have been well characterized^18-21^, the interaction and behavior of multiple phases within an open channel has been less extensively studied. Previously, in an open two-phase system driven by spontaneous capillary flow (SCF), we found that a single aqueous droplet within an open channel demonstrates different behavioral modes (e.g., translation, displacement, remaining stationary) largely governed by the interfacial tension between the droplet and the carrier phase, the contact angle of the droplet and carrier phase on the channel surface, and the velocity of the carrier phase.^17^ In the present manuscript, we use two of these behavioral modes (Figure S1) to create open channel manipulation modules driven by capillary flow: “shift mode”, in which an aqueous droplet wets all sides of the open channel and is translated downstream by the carrier phase (Figure S1i-ii), and “raft mode”, in which an aqueous droplet completely detaches from the channel and is displaced downstream by the carrier phase (Figure S1iii-iv).^17^ Shift mode occurs when an aqueous droplet in the channel precedes the advancing front of the carrier fluid, and the carrier fluid does not pass in front of the droplet; alternatively, raft mode occurs when the carrier fluid surrounds the droplet and simultaneously flows in front of and behind the droplet. Notably, in the case of both behavioral modes, the carrier fluid governs the overall dynamics of the system, as the pipetted droplets are entrained by the carrier fluid and transported downstream.

Within our open-channel platforms, we designed channel dimensions to fall within the flow regime governed by SCF^1,10^ to ensure capillary-driven flow, and incorporated a rounded channel geometry to negate flow along a wedge (i.e., Concus-Finn flow)^22^(Figure 1). Further, we fabricated our platform with poly (methyl methacrylate) (PMMA) to provide the desired wettability between droplets, the carrier phase, and the channel surface; specifically, with contact angles of 78° between PMMA and the aqueous droplet and ≈ 12.5° between PMMA and the organic carrier phase. Additionally, we enclosed our open channel platform within a humidified Omnitray™ (rectangular petri dish) surrounded by sacrificial water (1.5 mL in ≈ 50 μL droplets) to prevent evaporation of droplets from the channels during prolonged incubation times (i.e., hours). Our open channel system offers advantages to closed systems as we can add droplets directly to the channel with a pipette and initiate flow of the carrier phase through simple pipetting into the inlet reservoir (Figure 1).

### Open channel droplet incubation and transport

Inputting fluids into a typical microfluidic channel commonly requires dedicated ports and connectors. Addition of droplets into an open microfluidic channel, on the other hand, can be performed directly and at any location in the channel. Delivering small droplets (0.5-2.0 μL) with a pipette is a common approach available to most laboratories; alternatively, smaller droplets can be inputted through other approaches such as Acoustic Droplet Ejection methods.^23,24^ While depositing droplets on a surface is straightforward, removal or transfer of small droplets from a surface at subsequent time points is challenging; for example, pipetting is unreliable and tedious, as part of the droplet often remains behind on the surface. Digital microfluidics, also referred to as Electrowetting on Dielectric (EWOD), provides a method to move small droplets, but requires the use of electrical components.^25-27^ There is a need for simple systems in which droplets can be pipetted onto an unmodified surface, incubated *in situ* for a desired period, and then passively manipulated or transferred.

Previously, we demonstrated addition of droplets to an organic carrier phase as the carrier fluid was flowing downstream, enabling droplet transport but limiting the incubation time of the pipetted droplets to the time required to reach the outlet.^17^ Here, we present a different and adaptable platform where we place multiple discrete droplets into an open channel (in the absence of the organic carrier phase), incubate the droplets for a desired time, and then passively transport the droplets to a different location on chip *via* capillary-driven flow of an organic carrier phase (Figure 2). When multiple droplets are placed in series within an empty single open channel, translation of the droplets in shift mode leads to coalescence, as they merge with each subsequent droplet in the channel (Figure 2A). This functionality can be beneficial for analyses requiring pooling of multiple samples (e.g., discovery assays). For applications where droplet coalescence is not desired, we designed a separate flow path that we refer to as a ‘bypass channel’. The bypass channel enables the immiscible carrier fluid to separate each discrete droplet (thereby preventing coalescence) and transport the droplets downstream via raft mode (Figure 2B-C). When the carrier phase reaches the bypass, which is positioned upstream of the droplet, part of the flow of the carrier fluid diverts through the bypass, while the remainder of the carrier phase continues in the main channel; the diverted flow fills the space between each droplet, while the nondiverted flow continues to drive the droplets through the main channel (Figure 2B-C). Initially, we observed droplet disruption (e.g., droplet breakup and/or flow through the bypass) and an increased flow rate through the bypass, which resulted in stagnation of the carrier phase flow in the bends of the main channel and prevented droplets from reaching the outlet (Figure 2B). To mitigate this droplet disruption, we designed the bypass channels with a step (Figure 2C-D) to increase the hydrodynamic resistance (and therefore decrease the flux) through the bypass and maintain a sufficient flow rate in the main channel, ensuring that the droplets reached the outlet without breaking apart.

It is important to have a generalizable set of rules for designing bypass channels, to enable extension of our method to channels of different dimensions and geometries. We generated an analytical model that describes the ratio of fluid fluxes through the main channel relative to the bypass with respect to the fluidic resistance associated with each flow path. Deriving a model from a generalized Lucas-Washburn-Rideal law for open channels^18^ (SI), we found the relationship between the fluidic resistances and flux to be:

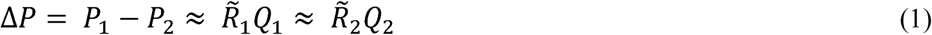

where *P* is the pressure drop across our bypass system (*P*_*1*_ and *P*_*2*_ refer to the pressure drops at the nodes before and after the bypass, respectively), *R*_*1*_ and *R*_*2*_ are the resistances in the main channel and bypass channel, respectively, and *Q*_*1*_ and *Q*_*2*_ are the flux through the main channel and the bypass channel, respectively (SI). The fluidic resistance of the capillary-driven flow in our system can be described by Equation 2:

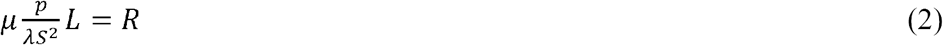

Where *S* is the cross-sectional area, *L* is the channel length, μ is the liquid viscosity, *p* is the total perimeter, and *λ* is the friction length.^28^ The resistance in our system inversely correlates to the cross-sectional area and the flux of fluid through the channel (SI). Solving for the ratio of the fluxes between the main channel and the bypass channel and then inserting the physical dimensions of our system into Equation 2 (as detailed in the SI) yielded a flux ratio of *Q*_*2*_*/Q*_*1*_ = 3.18 wherein the flux through the bypass channel (*Q*_*2*_) is greater than the flux through the main channel (*Q*_*1*_). However, through incorporation of a step, we are able to increase the resistance through the bypass channel and ensure less diversion of flow from the main channel, altering the flux ratio to *Q*_*2*_*/Q*_*1*_ = 2.68. This relation demonstrates that the resistance through the bypass can be manipulated by altering the geometry of the step (i.e., larger step, increased resistance and decreased flux), enabling adaptation for different geometries and channel lengths. With different manufacturing techniques, even greater ratios can be designed. Further, while the flux is still greater in the bypass than the main channel, the decrease in the flux through the bypass afforded by the step enabled sufficient flow in the main channel to drive droplets towards the outlet and prevent stagnation of the carrier fluid in the channel curves (Figure 2C). The derived model provides a framework for adapting the bypass system with different geometries and acts as a useful tool for quantification of flow and hydrodynamic resistance in open channels, as well as for guiding optimization of droplet flow in open channel systems.

The ability to transport small volumes with SCF lays the foundation for future open-channel platforms that integrate processes such as cell culture or biochemical reactions with small-volume readouts such as mass spectrometry and immunoassays. Overall, the bypass system is designed to incubate and manipulate droplets without extensive user interaction or difficult pipetting steps. Ease of use is demonstrated through a simple two-step process (droplet addition followed by carrier fluid addition) without the need for adjusting flow rates and flow directions for droplet manipulation. Due to the fabrication and droplet addition methods used, this technique is currently limited to low-throughput applications and droplet volumes compatible with micropipettes; however, the bypass platform can be expanded to include more droplets or to be compatible with arrays of different geometries.

### Controlled and adjustable droplet splitting in open channels

Building upon the droplet handling capabilities described for incubation and transport of multiple droplets, we demonstrate the ability to controllably split droplets within an open channel. Droplet splitting is an important feature that can extend the use of valuable or small volume samples by generating identical replicates and creating arrays for multiplexing. Using traditional droplet splitting geometries previously developed for closed-channel droplet-based microfluidics^7,8^, we designed T junctions to split incoming droplets, where a droplet entering the junction fills each branch of the junction until it is slowed down by an expansion in the geometry of the channel (Figure 3A)^29,30^; once the junction is filled, the droplet splits relative to the length of each branch (Figure 4A). By tuning the lengths of the left and right branches of the T junction, we can generate symmetric or asymmetric droplets (Figure 3B-C), enabling users to generate variable volume aliquots from an original sample simply by changing the device dimensions. Further, the splitting of symmetric and asymmetric droplets was reproducible between devices within an array (Figure 3D). Some variability was observed between multiple arrays due to manual micromilling artifacts, but inter-array variability can be alleviated through fabrication with more consistent methods such as injection molding and advanced automated milling.^11,14^

To demonstrate the workflow for a potential application of the open channel droplet splitting platform, we present a model experimental system for on-chip reagent delivery and reactions. We pipetted aqueous droplets tinted with yellow dye (to model primary reagents) upstream of a T junction and aqueous droplets tinted with green dye (to model secondary reagents) downstream of the T junction. We then loaded the inlet reservoir with carrier fluid to initiate flow. The carrier fluid flow drove the yellow droplet through the T junction, wherein the droplet split and its components were delivered to multiple samples (green droplets) downstream of the junction (Figure 4). The ability to preload the platform with reaction reagents allows users to generate multiplexed arrays for subsequent passive reagent delivery with minimal user handling; additionally, altering the channel distance and geometry after mixing can be used to adjust the incubation time of the reaction.^8^

## Conclusion

In this work, we develop essential features for immiscible droplet manipulation in capillary-driven open systems. Open channels offer several advantages over closed channels including pipette accessibility, manufacturability, customizability, and ease of use. Using these features, we created a generalized open channel platform for addition, incubation, and translation of multiple droplets and an open channel platform for droplet splitting and delivery. These platforms build upon prior work describing the fundamental behavior of single droplets in open biphasic systems by providing previously unavailable user functionalities (e.g., incubation of multiple droplets within a channel, droplet splitting) and creating foundational systems that can be customized and adapted for a range of experimental needs. Traditional closed-channel droplet microfluidics provides high throughput capabilities that can accommodate >10^7^ samples with droplet volumes reaching 10^−15^ L and can integrate with large-capacity screening instrumentation (LC/MS, high speed microscopy, etc.), greatly increasing the abilities of researchers to perform high throughput experimentation.^3^ Open-channel droplet microfluidics aims to address a different scale and set of experimental applications for researchers performing smaller screening studies (tens or hundreds of samples) with higher volumes (μL-nL) that do not require the extensive infrastructure nor cost associated with high throughput droplet microfluidics; further, we anticipate that our platform offers increased user accessibility, tractability, streamlining, and ease of use that allows for easy integration with existing experimental protocols and sample generation tools (e.g., pipettes, liquid handling robot). Future work with these platforms will include increasing the capacity of the bypass system for larger droplet arrays and expanding the droplet splitting capabilities to accommodate a wider range of splitting ratios. Further, in future investigations our platforms could be extended to smaller scales with the use of high resolution fabrication techniques and lower-volume liquid handlers or dispensers. In the future, we envision adaptation of these foundational platforms will enable users to expand and customize their current experimental toolbox for studies relating to drug screening, microscale reactions, and the “-omics” fields (e.g., metabolomics, proteomics).

## Supporting information

Open Channel Droplet-Based Microfluidics SI

Figure 1B Shift Mode

Figure 1C Raft Mode

Figure 2A Open Channel

Figure 2B Open Channel with Bypass

Figure 2C Open Channel with Stepped Bypass

Figure 3B Symmetrical Droplet Splitting

Figure 3C Asymmetrical Droplet Splitting

Figure 4 Droplet Splitting and Mixing

## Acknowledgments

This work was supported by the University of Washington and by the National Science Foundation Graduate Research Fellowship Program under Grant No. DGE-1256082. Any opinions, findings, and conclusions or recommendations expressed in this material are those of the author(s) and do not necessarily reflect the views of the National Science Foundation.

## References

1. Berthier, J.; Brakke, K.A.; Berthier, E. Open Microfluidics. Wiley, 2016.

2. Teh, S.Y.; Lin, R.; Hung, L.H.; Lee, A.P. Droplet microfluidics. Lab Chip, 2008, 8, 198–220.

3. Shang, L.; Cheng, Y.; Zhao, Y. Emerging Droplet Microfluidics. Chem. Rev., 2017, 117, 7964–8040.

4. Frenz, L.; Blank, K.; Brouzes, E.; Griffiths, A.D. Reliable microfluidic on-chip incubation of droplets in delay-lines. Lab Chip, 2009, 9, 10, 1344–1348.

5. Boukellal, H.; Selimovic, S.; Jia, Y.; Cristobal, G.; Fraden, S. Simple, robust storage of drops and fluids in a microfluidic device. Lab Chip, 2009, 9, 331–338.

6. Abate, A.R.; Hung, T.; Mary, P.; Agresti, J.J.; Weitz, D.A. High-throughput injection with microfluidics using picoinjectors. PNAS, 2010, 107, 45, 19163–19166.

7. Link, D.R.; Anna, S.L.; Weitz, D.A; Stone, H.A. Geometrically mediated breakup of drops in microfluidic device. Phys. Rev. Lett., 2004, 92, 054503.

8. Song, H.; Chen, D.L.; Ismagilov, R.F. Reactions in droplets in microfluidic channels. Angew. Chem. Int. Ed. Engl., 2006, 45, 44, 7336–7356.

9. Li, C.; Boban, M.; Tuteja, A. Open-channel, water-in-oil emulsification in paper-based microfluidic devices. Lab Chip, 2017, 1436.

10. Casavant, B.P.; Berthier, E.; Theberge, A.B.; Berthier, J.; Montanez-Sauri, S.I.; Bischel, L.L.; et al. Suspended Microfluidics. PNAS, 2016, 110, 10111–10116.

11. Guckenberger, D.J.; de Groot, T.E.; Wan, A.M.D.; Beebe, D.J.; Young, E.W.K. Micromilling: A method for ultra-rapid prototyping of plastic microfluidic devices. Lab Chip, 2015, 15, 11, 2364–2378.

12. de Groot, T.E.; Veserat, K.S.; Berthier, E.; Beebe, D.J.; Theberge, A.B. Surface-tension drive open microfluidic platform for hanging droplet culture. Lab Chip, 2016, 16, 334.

13. Barkal, L.J.; Theberge, A.B.; Guo, C.J.; Spraker, J.; Rappert, L.; Berthier, J.; et al. Microbial metabolomics in open microscale platforms. Nature Communications, 2016, 7, 10610.

14. Lee, U.N.; Su, X.; Guckenberger, D.J., Dostie, A.M.; Zhang, T.; Berthier, E.; et al. Fundamentals of rapid injection molding for microfluidic cell-based assays. Lab Chip, 2018, 18, 496–504.

15. Lee, Y.; Choi, J.W.; Yu, J.; Park, D.; Ha, J.; Son, K.; et al. Microfluidics within a well: an injection-molded plastic array 3D culture platform. Lab Chip, 2018, Advance Article.

16. Huemmer, D.; Bachler, S.; Kohler, M.; Blank, L.M.; Zenobi, R.; Dittrich, P.S.; Microfluidic platform for multimodal analysis of enzyme secretion in nanoliter droplet arrays. Anal. Chem., 2018, xxxxx

17. Lee, J.J.; Berthier, J.; Brakke, K.A.; Dostie, A.M.; Theberge, A.B.; Berthier, E. Droplet behavior in Open Biphasic Microfluidics. Langmuir, 2018, 34, 18, 5358–5366.

18. Berthier, J.; Gosselin, D.; Berthier, E. A generalization of the Lucas-Washburn-Rideal law to composite microchannels of arbitrary cross section. Microfluid. Nanofluid., 2015, 19, 3, 497–507.

19. Chen, Y.F.; Tseng, F.G.; Chein, S.Y.C.; Chen, M.H.; Yu, R.J.; Chieng, C.C. Surface tension driven flow for open microchannels with different turning angles. Microfluid Nanofluid, 2008, 5, 193–203.

20. Lade Jr., R.K.; Jochem, K.S.; Macosko, C.W.; Francis, L.F. Capillary Coatings: Flow and Drying Dynamics in Open Microchannels. Langmuir, 2018, 34, 7624–7639.

21. Yang, D.; Krasowska, M.; Priest, C.; Popescu, M.N.; Ralston, J. Dynamics of capillary-driven flow in open microchannels. J. Phys. Chem. C., 2011, 115, 18761–18769.

22. Concus, P.; Finn, R. On the behavior of a capillary surface in a wedge. PNAS, 1969, 2, 63, 292–299.

23. Hadimioglu, B.; Stearns, R.; Ellson, R. Moving Liquids with Sound: The Physics of Acoustic Droplet Ejection for Robust Laboratory Automation in Life Sciences. Journal of Laboratory Automation, 2016, 21, 1, 4–18.

24. Demirci, U. Acoustic Picoliter Droplets for Emerging Applications in Semiconductor Industry and Biotechnology. Journal of Microelectromechanical Systems, 2006, 15, 4, 957–966.

25. Choi, K.; Ng, A.H.C.; Fobel, R.; Wheeler, A.R. Digital Microfluidics. Annu. Rev. Anal. Chem., 2012, 5, 413–440.

26. Ng, A.H.C.; Li, B.B.; Chamberlain, M.D.; Wheeler, A.R. Digital Microfluidic Cell Culture. Annu. Rev. Biomed. Eng., 2015, 17, 91–112.

27. Jones, T.B. On the relationship of dielectrophoresis and electrowetting. Langmuir, 2002, 18, 11, 4437–4443.

28. Lee, J.J.; Karampelas, I.H.; Brakke, K.A.; Theberge, A.B.; Berthier, E.; Berthier, J. Capillary flow of solvents and aqueous liquids in rounded open microgrooves. Microfluid. Nanofluid. Submitted.

29. Mehrabian, H.; Gao, P.; J.J. Feng, J.J. Wicking flow through microchannels. Physics of Fluids, 2011, 23, 122108.

30. Berthier, J; Brakke, K.A.; Gosselin, D.; Navarro, F.; Belgacem, N.; Chaussy, D.; Berthier, E. On the halt of spontaneous capillary flows in diverging open channels. Med. Eng. Phys. 2017, 75–80.

