## Supplementary material for "Open channel droplet-based microfluidics": Open Channel Droplet-Based Microfluidics SI

The supporting information for “Open Channel Droplet-Based Microfluidics” includes additional information that readers might find useful for adapting the open channel systems presented here for their own work; this includes detailed schematics and dimensions of each platform described in this work and corresponding computer aided design (CAD) files, a derivation of the analytical model used to describe flow in the bypass system, an illustrated workflow of the droplet length measuring protocol used for Figure 3D, and videos showing the use and function of each platform in this work.

Supporting information contents:

S1: Schematics and images of droplet behavior modes

S2: Schematics and drawings detailing specific device dimensions

S3: Derivation and calculation of analytical model describing resistance and flux in the bypass platform

S4: Illustrated workflow and description of droplet length measuring protocol

Additional supporting information included:

S5: Table of video files (.mpg) showing platform use and function for device type

S6: Table of Solidworks (.zip/.STEP) CAD files for each device

**S1: Droplet Behavior Modes in Open Channel Platform**
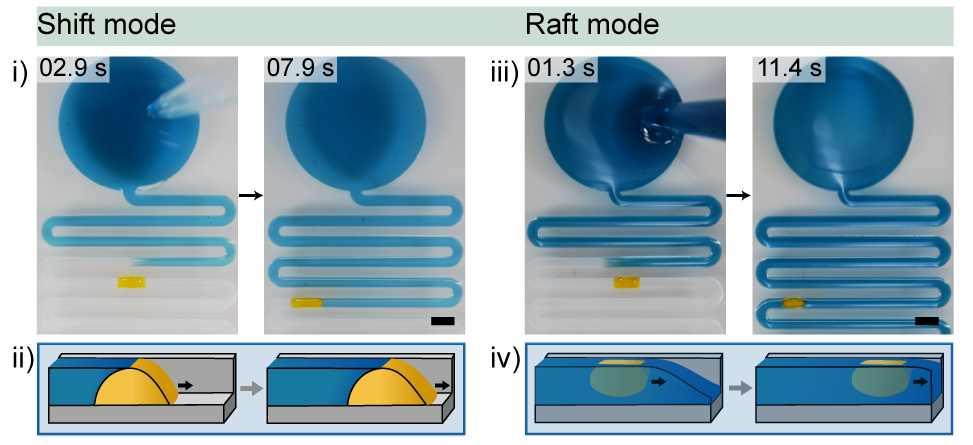

Figure S1: A single static aqueous droplet (yellow) is pipetted into the channel and can display either shift mode ((i) and (ii)) or raft mode ((iii) and (iv)) as carrier fluid (blue) moves the aqueous droplet down channel.^1^ For shift mode, the carrier phase is 1-pentanol; for raft mode, the carrier phase is toluene. Scale bar: 2 mm. Schematics in (ii) and (iv) are reproduced from Lee et al.^1^ Videos for (i) and (iii) are included (Videos S1 and S2).

Reference:

1. Lee, J.J.; Berthier, J.; Brakke, K.A.; Dostie, A.M.; Theberge, A.B.; Berthier, E. Droplet behavior in Open Biphasic Microfluidics. *Langmuir*, **2018**, 34, 18, 5358-5366.

**S2: Detailed Device Schematics and Dimensions**

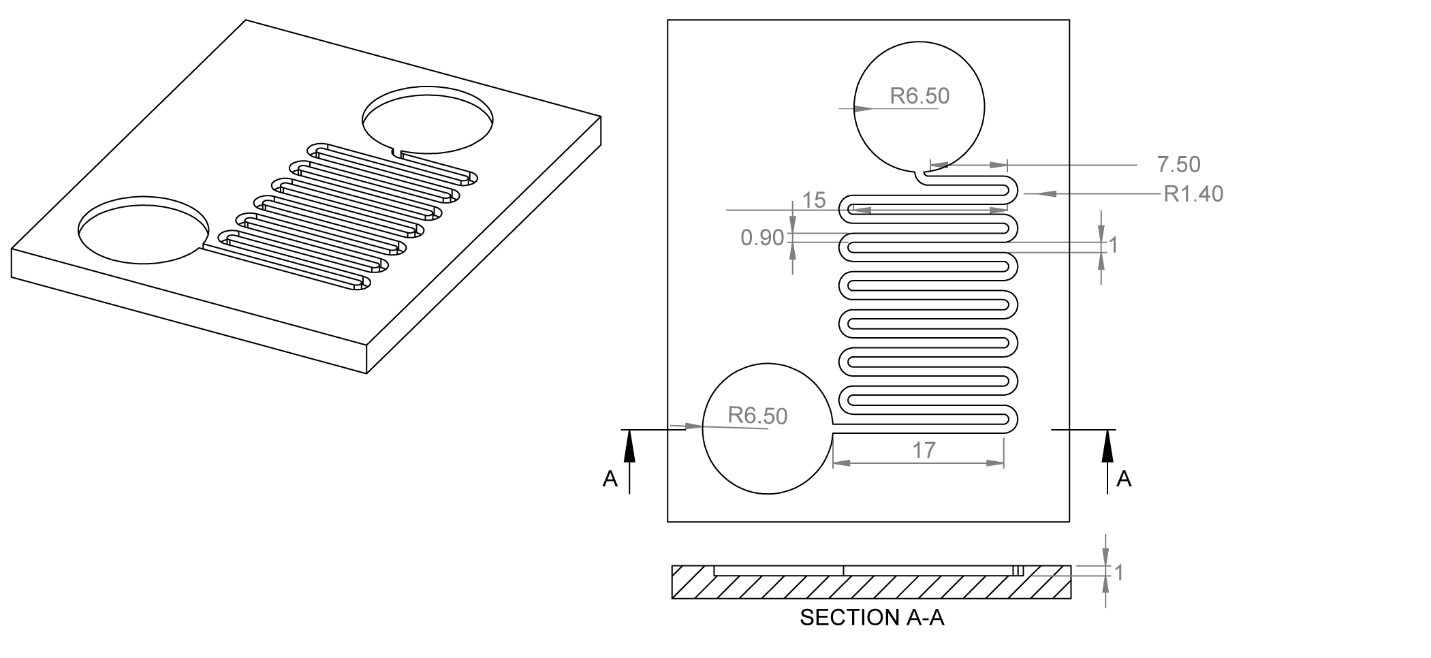

Figure S2i: Detailed device schematics illustrating device dimensions for the Coalescence Device (Figure 2A). Units in mm.

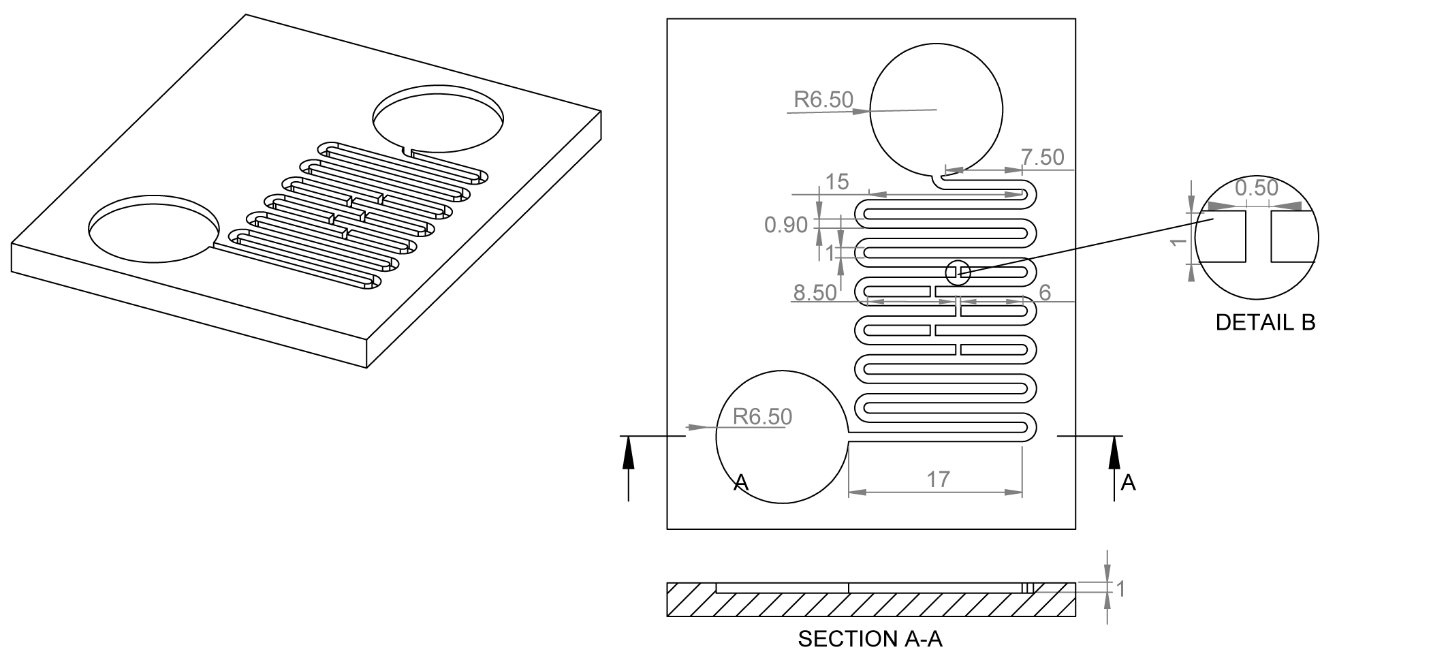

Figure S2ii: Detailed device schematics illustrating device dimensions for the Bypass Device without a step (Figure 2B). Units in mm.

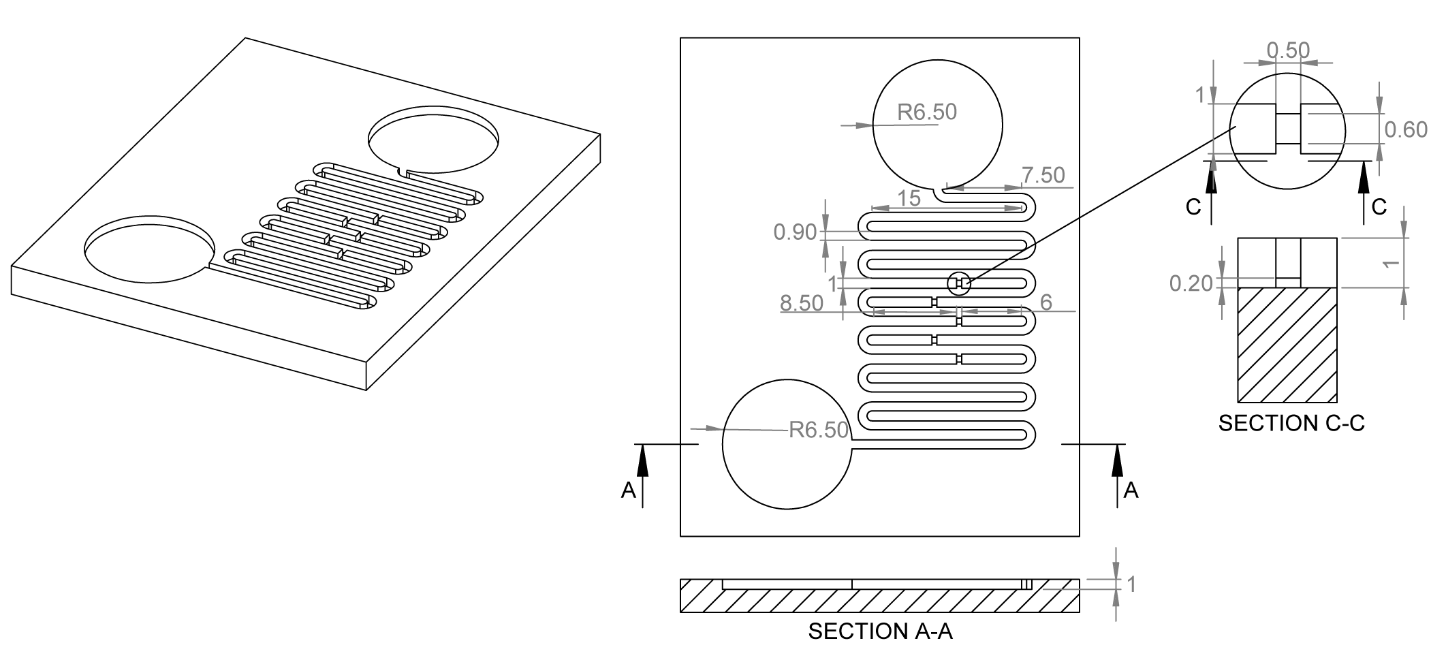

Figure S2iii: Detailed device schematics illustrating device dimensions for the Bypass Device with a step (Figure 2C). Units in mm.

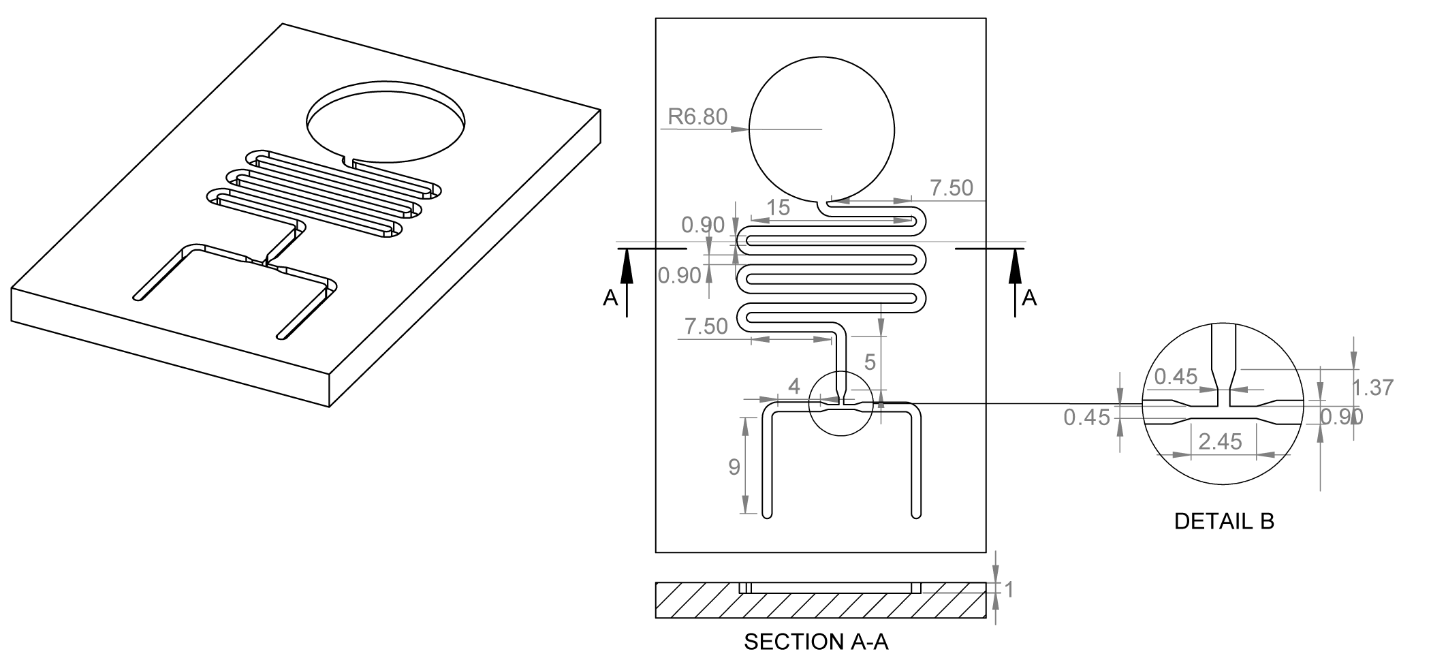

Figure S2iv: Detailed device schematics illustrating device dimensions for the symmetric splitting T junction device (Figure 4B). Units in mm.

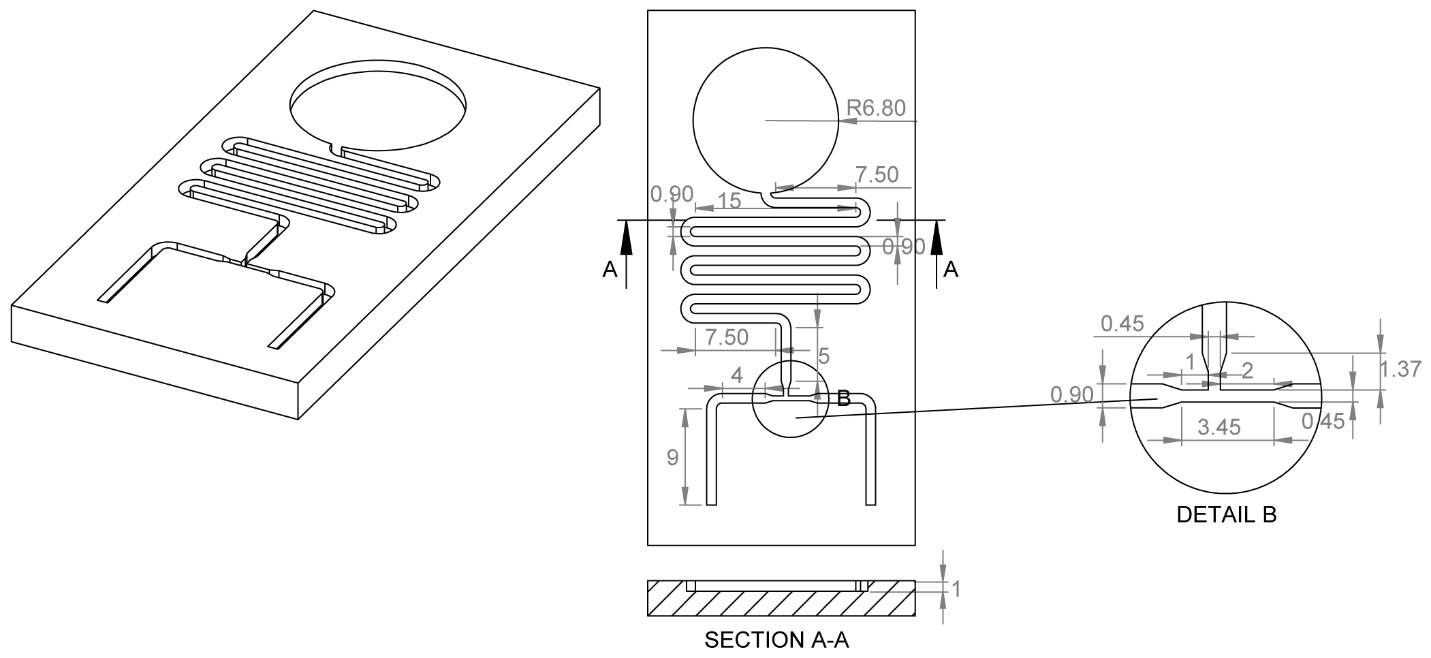

Figure S2v: Detailed device schematics illustrating device dimensions for the asymmetric splitting T junction device (Figure 4C). Units in mm.

**S3: Derivation of Analytical Model for Resistance and Flux**

In this model, we considered an open channel system with a bifurcation and two nodes, where the flows partition at the first node (green dot) and join at the second (red dot). Between the two nodes are two different branches (blue and yellow outlines); Branch 1 is an extension of the main channel, and Branch 2 is the bypass channel with a step in the middle (Figure S2i).

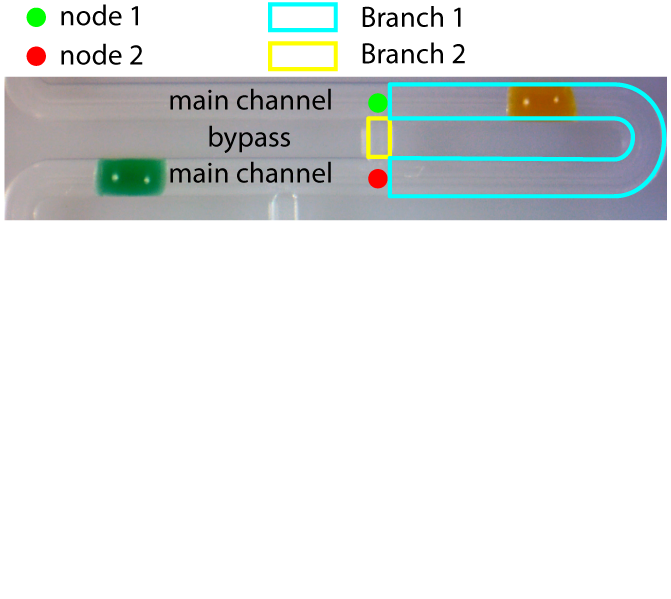

Figure S2i: Labeled top view image of open channel system.

Based on previous work^1^ on the creation of a generalized Lucas-Washburn law for varying shapes driven by capillary flow and application of the analogy of an electric circuit, we can write:

$\Delta P= P_{1}-P_{2}\approx\tilde{R}_{1}Q_{1}\approx\tilde{R}_{2}Q_{2}$	(1)

$\mu\frac{p}{\lambda S^{2}}L=R$	(2)

Where P_1_ and P_2_ are the pressures at node 1 and node 2, respectively, R is the resistance, L is the length, Q is the flux, S is the channel cross sectional area, μ is the liquid viscosity, p is the total perimeter, and 𝜆 is the friction length.^2^

Solving equation 1 for Q:

$\left( \frac{\tilde{R}_{1}}{\tilde{R}_{2}} \right)=\left( \frac{Q_{2}}{Q_{1}} \right)$	(3)

From Equation 3, we have a relation that describes the flow rate in terms of the resistance and can be used to calculate the specific resistance for each channel with physical parameters from the system. However, Branch 2 has a heterogenous cross section due to the step; therefore, the resistances of each section of the bypass channel must be considered to find the total resistance through Branch 2:

$\left( \frac{\tilde{R}_{1}}{\tilde{R}_{2}} \right)=\left( \frac{\tilde{R_{1}}}{\tilde{R_{i}}+\tilde{R_{ii}}+\tilde{R_{iii}}} \right)=\left( \frac{Q_{2}}{Q_{1}} \right)$	(4a)

And plugging in Equation 2:

$\left( \frac{\mu\frac{p_{1}}{\lambda_{1}S_{1}^{2}}L_{1}}{\mu\frac{p_{i}}{\lambda_{i}S_{i}^{2}}L_{i}+\mu\frac{p_{ii}}{\lambda_{ii}S_{ii}^{2}}L_{ii}+\mu\frac{p_{iii}}{\lambda_{iii}S_{iii}^{2}}L_{iii}} \right)=\left( \frac{Q_{2}}{Q_{1}} \right)$	(4b)

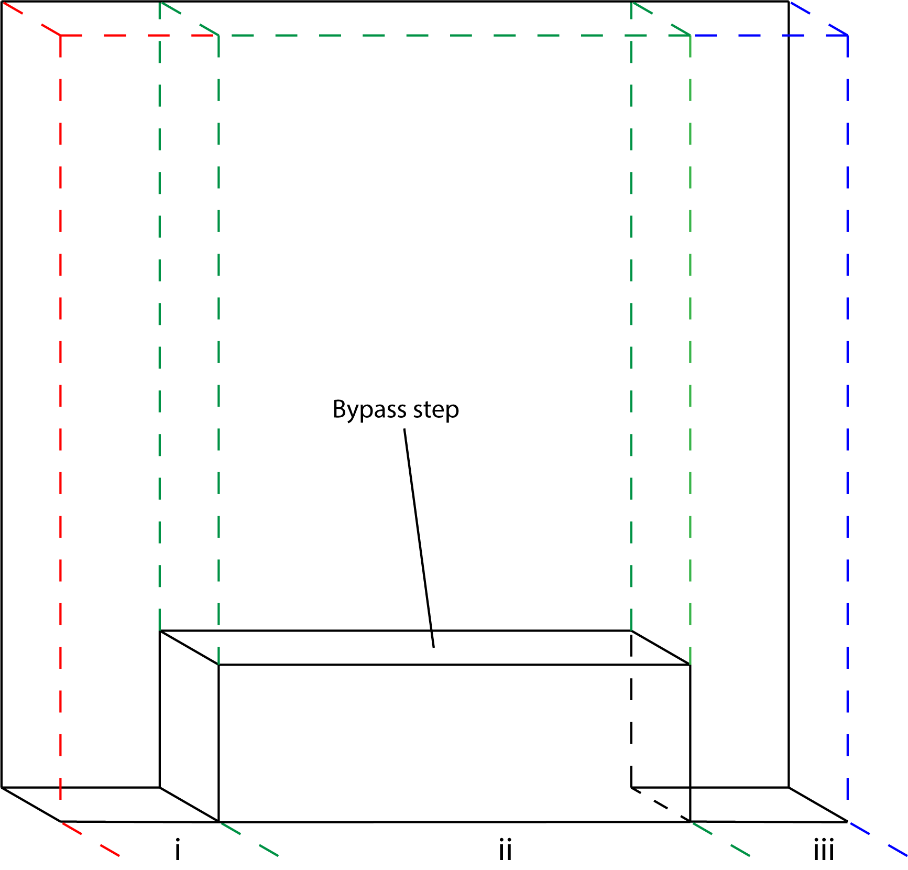

Figure S2ii: Isometric view of cross section of Branch 2 (Bypass)

Where i, ii, and iii represent the separate sections within the bypass channel (Figure S2ii). Plugging in the physical parameters into Equation 4b provides the ratio of the fluxes between the bypass channel and the main channel. For our specific stepped bypass platform:

$\frac{Q_{2}}{Q_{1}}=2.68$	(5a)

And without a step in the bypass channel:

$\frac{Q_{2}}{Q_{1}}=3.18$	(5b)

These calculations demonstrate that incorporation of the step into the bypass decreases the flux through the bypass by increasing the resistance through the bypass channel. The decreased flux through the bypass (Branch 2) corresponds to an increased flux through the main channel (Branch 1), which drives the droplets completely through the curves in the channels and allows them to flow to the outlet.

References:

2. Berthier, J.; Gosselin, D.; Berthier, E. A generalization of the Lucas-Washburn Rideal law to composite microchannels of arbitrary cross section. Microfluid. Nanofluid., 2015, 19, 3, 497-507.
3. Lee, J.J.; Karampelas, I.H.; Brakke, K.A.; Theberge, A.B.; Berthier, E.; Berthier, J. Capillary flow of solvents and aqueous liquids in rounded open microgrooves. Microfluid. Nanofluid. Submitted.

**S4: Workflow for Droplet Length Measurements**

To measure the length of the aqueous droplets in our channels, we designed and fabricated T junction platforms without outlet reservoirs. As with the devices with the outlet reservoir, we pipetted a 3 μL droplet upstream of the T junction and the added carrier fluid to the inlet reservoir. After initiation of spontaneous capillary flow (SCF) and splitting of the droplet, daughter droplets flowed to the end of the channel, where they stopped and were compressed as the device filled with carrier fluid. Once the device was completely filled and flow had completely ceased, the daughter droplets at the end of the channel were imaged. Images of droplets were then analyzed with ImageJ. Specifically, A-B) images were opened (symmetric droplet image) and scaled (Analyze, set scale); C) the “Segmented Line” tool was then selected and a vertical line was drawn from the top of each droplet to the bottom. D) The segmented line was measured (Analyze, measure) and droplet lengths were recorded.

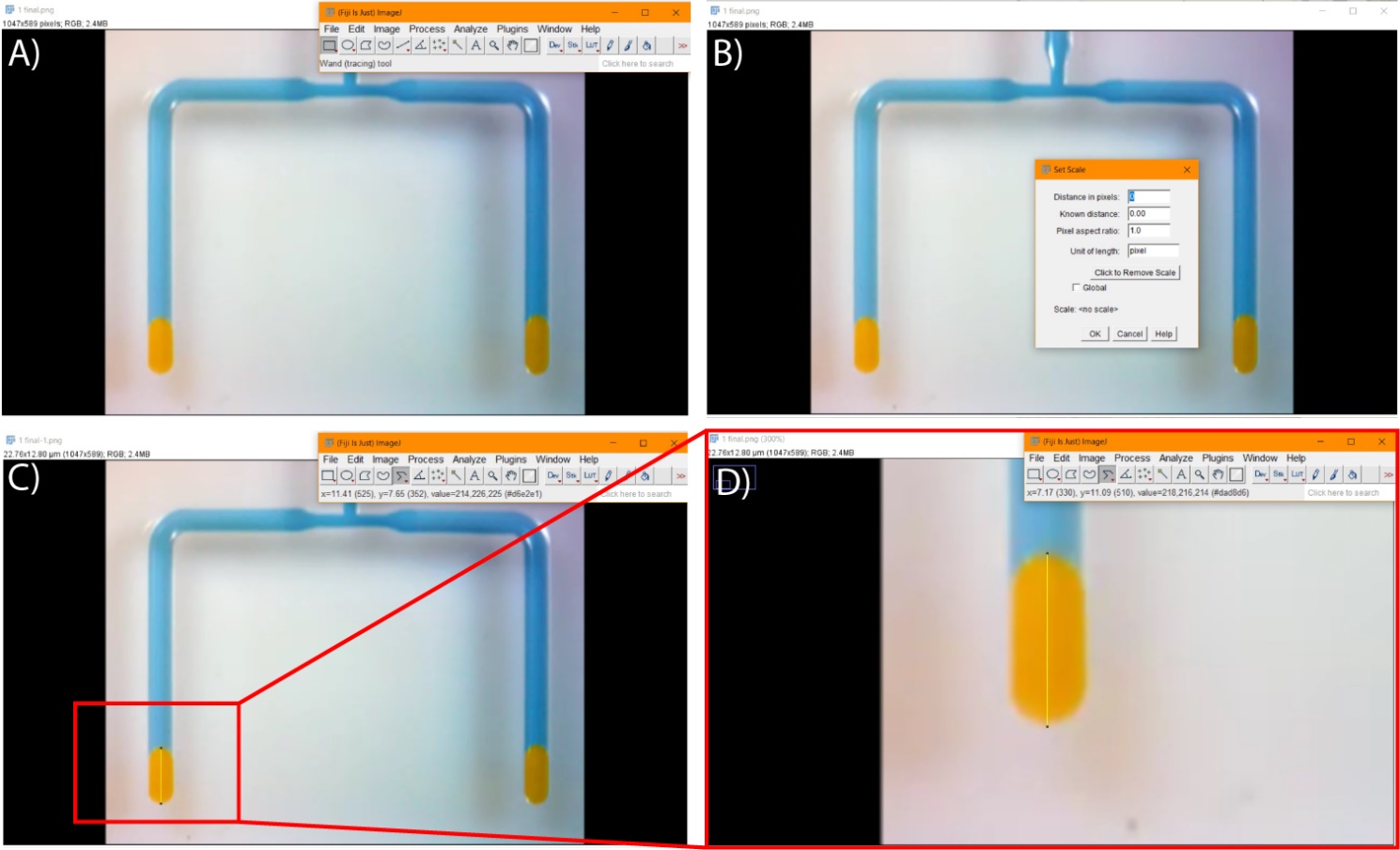

Figure S3: Workflow for droplet length measurement. A) The image to be measured is opened using ImageJ; B) to set the scale, we selected “Analyze, set scale” and changed the scale from pixels to μm; C) the Segmented Line tool is selected, and a line is draw from one end of the droplet to the other (interface of droplet and carrier phase and boundary of droplet and end of channel); D) zoomed-in image illustrating the line drawn on the droplet.

**S5: Table of Video Files Included in Supporting Information**

| Video | File Name |
| --- | --- |
| S1 | Fig 1B Shift mode.mpg |
| S2 | Fig 1C Raft mode.mpg |
| S3 | Fig 2A Open channel.mpg |
| S4 | Fig 2B Open channel with bypass.mpg |
| S5 | Fig 2C Open channel with stepped bypass.mpg |
| S6 | Fig 3B Symmetric droplet splitting.mpg |
| S7 | Fig 3C Asymmetric droplet splitting.mpg |
| S8 | Fig 4 Droplet splitting and mixing.mpg |

**S6: Table of Design Files Included in Supporting Information**

| Device Figure | File Name |
| --- | --- |
| Figure 1B | Open channel.STEP |
| Figure 2A | Open channel.STEP |
| Figure 2B | Open channel with bypass.STEP |
| Figure 2C | Open channel with stepped bypass.STEP |
| Figure 3B | T junction 1-1.STEP |
| Figure 3C | T junction 1-2.STEP |
| Figure 4 | T junction 1-1.STEP |
